# Genome-centric portrait of the microbes’ cellulolytic competency

**DOI:** 10.1101/2020.03.09.984823

**Authors:** Yubo Wang, Liguan Li, Yu Xia, Feng Ju, Tong Zhang

## Abstract

Neither the abundance of the exo/endoglucase GH modules nor the taxonomy affiliation is informative enough in inferring whether a genome is of a potential cellulolytic microbe or not. By interpreting the complete genomes of 2642 microbe strains whose phenotypes have been well documented, we are trying to reveal a more reliable genotype and phenotype correlation on the specific function niche of cellulose hydrolysis. By incorporating into the annotation approach an automatic recognition of the potential synergy machineries, a more reliable prediction on the corresponding microbes’ cellulolytic competency could be achieved. The potential cellulose hydrolyzing microbes could be categorized into 5 groups according to the varying synergy machineries among the carbohydrate active modules/genes annotated. Results of the meta-analysis on the 2642 genomes revealed that some cellulosome gene clusters were in lack of the surface layer homology module (SLH) and microbe strains annotated with such cellulosome gene clusters were not certainly cellulolytic. Hypothesized in this study was that cellulosome-independent genes harboring both the SLH module and the cellulose-binding carbohydrate binding module (CBM) were likely an alternative gene apparatus initiating the formation of the cellulose-enzyme-microbe (CEM) complexes; and their role is important especially for the cellulolytic anaerobes without cellulosome gene clusters.

**Importance:** In the genome-centric prediction on the corresponding microbes’ cellulolytic activity, recognition of the synergy machineries that include but are not limited to the cellulosome gene clusters is equally important as the annotation of the individual carbohydrate active modules or genes. This is the first time that a pipeline was developed for an automatic recognition of the synergy among the carbohydrate active units annotated. With promising resolution and reliability, this pipeline should be a good add to the bioinformatic tools for the genome-centric interpretations on the specific function niche of cellulose hydrolysis.

## Background

In the era of high-throughput sequencing, the genetic information that is inherently whispering hints of the microbes’ function niches is becoming easily accessible (1, 2). However, the bottleneck remains largely on properly identifying and characterizing these genetic hints and inferring the microbes’ function potentials. In this study, we focus on the genome-centric interpretation on the specific function niche of cellulose hydrolysis. Traditional approaches, including the microscope observation, cultivation of the cellulose-degrading microbes, as well as purification and characterization of the cellulolytic enzymes (3, 4), have set a good foundation in understanding how the microbes and their enzymes may interact with the cellulosic substrates. Although it is believed that most of the cellulolytic microbes may still be hiding in plain sight due to the isolation bottleneck, access to their genome information has opened a new window to shed light on them.

Regarding to the genome-centric interpretations on the function niche of cellulose hydrolysis, current annotation approaches focus on tapping the diversity and the abundance of the individual carbohydrate active enzyme (CAZy) modules annotated. Applying the HMMsearch-based dbCAN annotation platform, referring to the well-curated CAZy database (5, 6), a decent amount of information on the abundances of the diverse CAZy modules in a genome could be obtained. However, often encountered in practice was a lack of confidence in predicting the microbes’ real cellulolytic competency based solely on the abundances of the relevant CAZy modules annotated. For example, a total number of 21 exo/endoglucanase GH modules in the genome of *Actinoplanes missouriensis* 431 could not point to a conclusion that this strain was able to hydrolyze cellulose (7); and it remains a puzzle why *Clostridium acetobutylicum*, with the cellulosome gene cluster identified in its genome, do not have the cellulose degrading capability (8–10).

What is in lack in current genome-centric interpretations is the recognition of the synergy among the individual CAZy modules and among the carbohydrate active genes harboring these CAZy modules, although such synergy is one of the highly-appreciated features in efficient cellulose hydrolysis (11, 12). Cellulolytic enzymes are known as modular proteins, the most straightforward synergy would occur among the diverse CAZy modules in one single gene/enzyme; e.g., if one gene has both the cellulose-binding module CBM6 and the exoglucanase module GH9, the CBM6 could help bring this GH9 to its action site. A higher level of synergy would occur among the diverse carbohydrate active enzymes in one microbe, on which aspect, cellulosomes is the most highly recognized synergy machinery in anaerobes; and the CEM complex initiated by hypha penetration is the more commonly observed synergy mechanism in aerobic cellulolytic *Fungi* (11). Although the carbohydrate active enzymes did not assemble into one entity as those in the cellulosome complexes (9, 17), in the CEM complex of some aerobic Fungi, physical closeness among the individual carbohydrate active enzymes sandwiched in between the Fungus cells and the cellulose substrates makes the synergy among these enzymes possible (18–22).

May the formation of the cellulosome independent CEM complexes be possible in anaerobes? This question is raised in the context of the fact that the number of the cellulolytic anaerobes is much larger than the number of anaerobes with cellulosome gene clusters. Most cellulolytic species were of their optimal growth rates when they adhere to the cellulosic substrate, and the microbe-cellulose contact is important for the host microbes to get easy access to the enzymatic hydrolyzing products (15, 16). It has also been reported that the excreted free cellulase would contribute little to the microbes’ cellulolytic activity (11, 15). Taken together, the physical closeness in the form of the CEM complex might be critical for microbial cellulose hydrolysis. It is not common to observe in anaerobes the physical apparatus like hypha to facilitate the physical penetration as in Fungus, having been reported in literature was that the hypothesized glycocalyx mediated microbe-cellulose contact in anaerobes (23). One of the objectives of this study is, by investigating complete genomes of the 2642 microbe strains whose phenotypes have been well characterized, to uncover potential alternative genetic machineries (in lieu of the cellulosome complexes) that may initiate the CEM complex formation through the microbe-cellulose adhesion, especially in anaerobes.

Physical link or physical closeness is important for the synergy interactions among the carbohydrate active units (13). One recent progress in the recognition of the physical-link among the carbohydrate active genes was the establishment of the polysaccharide-utilization loci (PUL) database (14). In this study, we are trying to introduce the annotation of two more features regarding the physical connections among the CAZy modules and among the carbohydrate active enzymes: 1) clustering patterns among the CAZy modules along genes; and 2) machineries that may facilitate the assembly or physical aggregation of the diverse carbohydrate active enzymes in one microbe.

To summarize, for a reliable genotype and phenotype correlation on the specific function niche of cellulose hydrolysis, starting with the meta-analysis of the complete genomes of the 2642 microbe strains, we are aiming to test the possibility of developing an annotation pipeline for: 1) an automatic recognition of the clustering patterns among the CAZy modules in carbohydrate active genes in genomes, 2) recognition of potential alternative genetic machineries for the CEM complex formation in microbes, and 3) categorization of the genomes of potential cellulolytic microbes. The applicability of the pipeline in the annotation of metagenome assembled genomes (MAGs) could be further tested with the annotation of 7904 reference genomes downloaded from NCBI.

## Results

### Co-occurring patterns among the CAZy modules in the carbohydrate active genes

Genes are the basic units encoding enzymes, presented in Figure 1 and the Appendix file 1 are the frequencies at which CAZy modules co-occurring with each other in same genes; and these frequencies were calculated from the CAZy modules in carbohydrate active genes annotated in the 2642 complete genomes.

**Figure 1.**
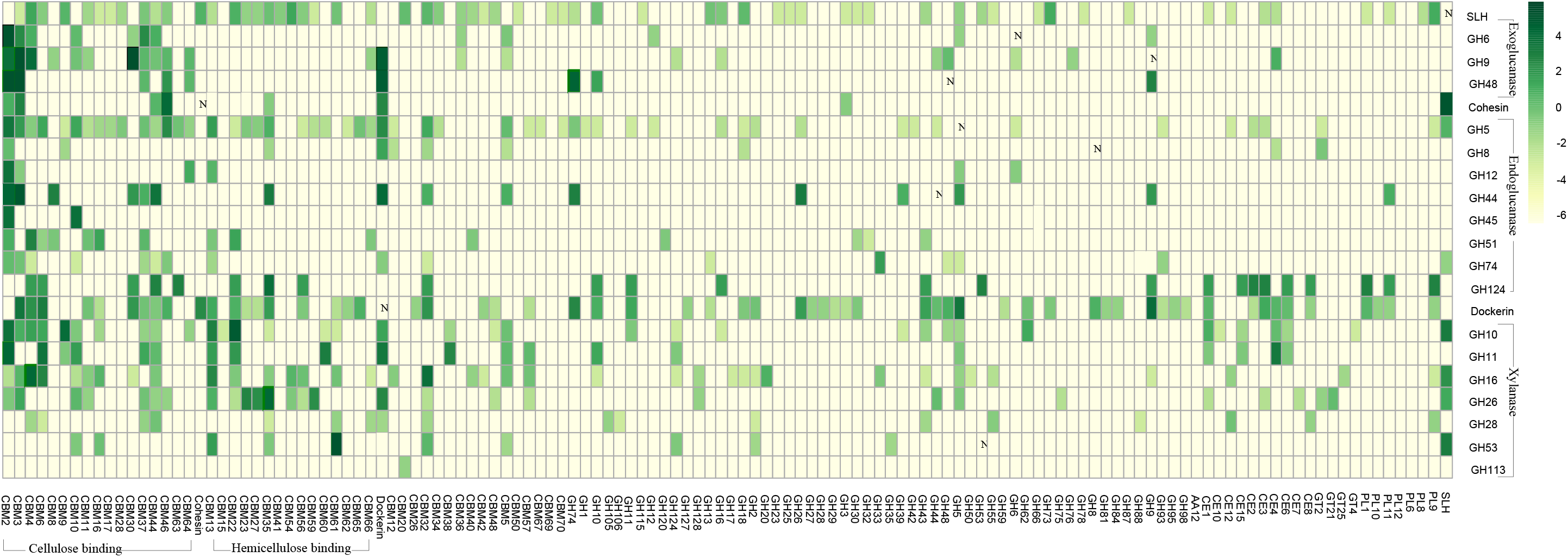
Co-occurrence frequencies among the CAZy modules Note: Vertically listed were the twenty-one selected CAZy modules, including the three exoglucanase GH modules, the eight endoglucanase GH modules, the seven xylanase GH modules, the cohesin, the dockerin and the SLH module. The CAZy modules lining horizontally were those modules being observed in same genes with at least one of the vertically listed CAZy modules. The scale bar on the right presented the co-occurring frequencies (x) in the log format of lg(x+0.01), and the plain number ‘x’ was summarized in the Appendix file 2.

One of the most distinctive co-occurrences was observed between the exoglucanase GH modules (GH6, GH9 and GH48) and the cellulose binding CBMs modules (dominantly CBM2, CBM3 and CBM30). Among the CAZy modules annotated in the 2642 complete genomes, 51% of the GH48 modules were observed being present in same genes with the CBM2 module; and CBM2 was also observed in 29% of the genes harboring the GH6 module. This is in accord with the reported importance of the CBM modules in: 1) the initiation of the exo/endoglucanase GH modules’ hydrolytic activity and 2) the progressiveness of the exoglucanase along the cellulose chains (11). Similarly observed was the co-occurrence between the xylanase GH modules (e.g., GH53, GH10) and the hemicellulose binding CBM modules (e.g., CBM61, CBM22), e.g., 26% of the genes harboring the GH10 module would also carry the CBM22 module.

Besides their high frequencies co-occurring with the cellulose-binding CBM modules, GH9 and GH48 were also the two modules with the highest frequencies co-occurring with the dockerin module, e.g. ~23% of the GH9-harboring genes were also identified with the dockerin module; and this suggested that GH9 and GH48 might be the two most common catalytic components in the cellulosome complexes. Collaboration between the exoglucanase and the endoglucanase was another important synergy pattern in cellulose hydrolysis; and this corresponded with the observation that ~20% of the exoglucanase GH48 module coexisted in same genes as the endoglucanase GH74 module.

### Categorization of the carbohydrate active genes

Part of the visualization of the CAZy module arrangement along carbohydrate active genes is demonstrated in Figure 2. According to the CAZy modules they harbor, the carbohydrate active genes could be classified into two broad categories: genes of the cellulosome gene clusters and carbohydrate active genes independent of the cellulosome gene clusters. As is summarized in Table 1, the cellulosome gene clusters consist of two parts, the scaffold genes (A1, A2 and A3) and genes of the catalytic components (A-s: dockerin + GH/CBM). The scaffold genes in the cellulosome gene clusters could be further categorized into three types (‘A1’, ‘A2’ and ‘A3’), according to whether the SLH module is initially in (type ‘A1’) or at least could be incorporated (type ‘A2’) into these scaffold genes. The integration of the ‘A2-a’ gene and the ‘A2-b’ gene by the dockerin and cohesion modules would incorporate the SLH module into the type ‘A2’ scaffolds. The scaffold genes of type ‘A3’ are in lack of the SLH module.

**Figure 2.**
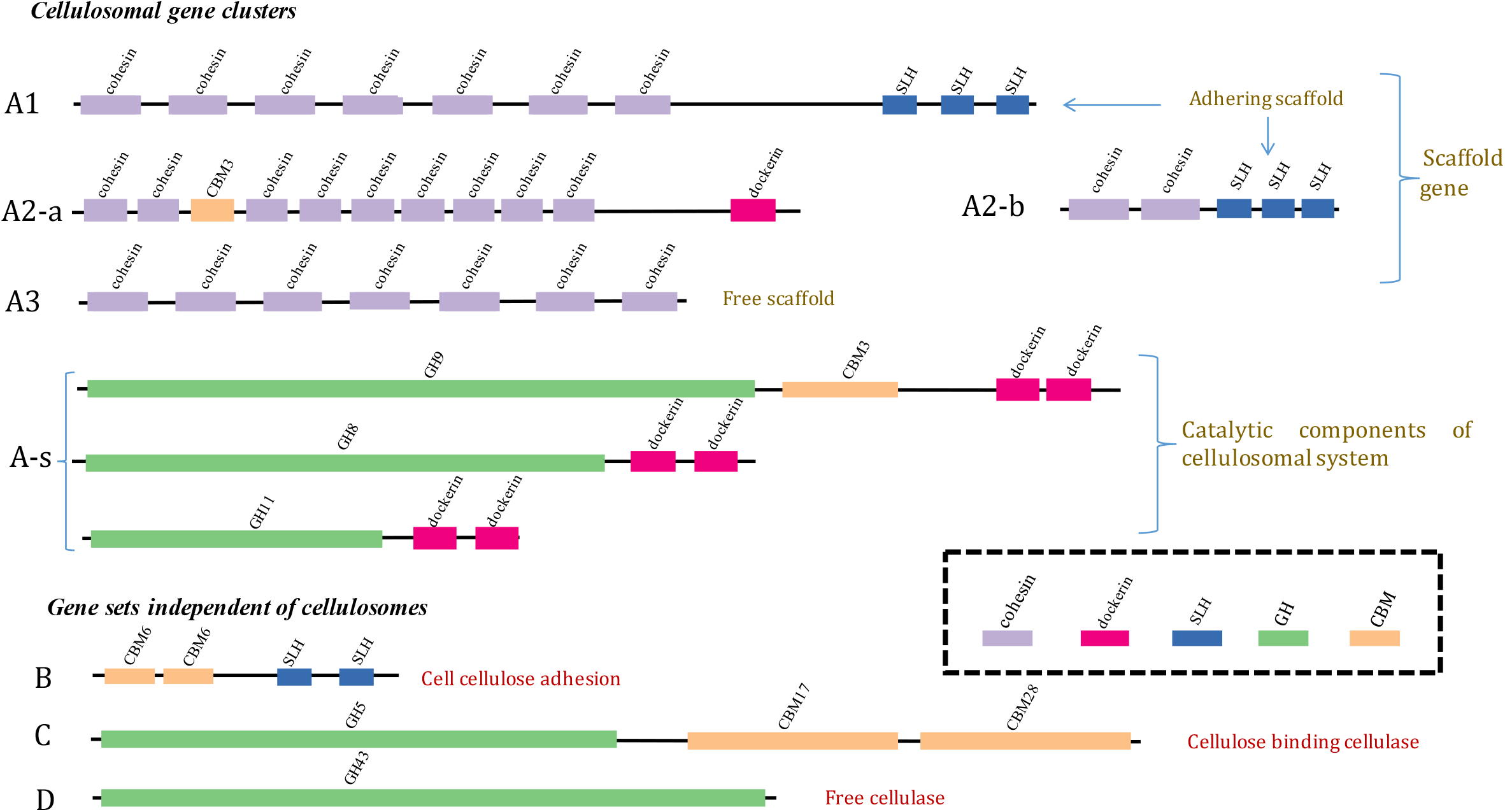
Illustration on the clustering patterns of the CAZy modules along gene

**Table 1.** Categorization of genes based on the CAZy modules they harbor

| Gene type |  |  | Abundance of modules in one gene |  |  |  |  | Gene Description |
| --- | --- | --- | --- | --- | --- | --- | --- | --- |
|  |  |  | SLH | Dokerin | Cohesin | GH | CBM |  |
| Cellulosomal<br>gene clusters | Scaffold<br>gene | A1 | >=1 | 0 | >=3 | >=0 | >=0 | Adhering scaffold<br>(Cohesin+Cohesin+...+Cohesin+SLH) |
|  |  | A2-a | >=0 | >=1 | >=3 | >=0 | >=0 | Adhering scaffold<br>(Cohesin+Cohesin+...+Cohesin+Dockerin) |
|  |  | A2-b | >=1 | >=0 | >=1 | >=0 | >=0 | Counterparts of A2-a<br>(Cohesin+SLH) |
|  |  | A3 | 0 | 0 | >=3 | >=0 | >=0 | Free scaffold<br>(Cohesin+Cohesin+...+Cohesin) |
|  | Catalytic<br>constituents | A-s | >=0 | >=1 | >=0 | >=1 | >=0 | Catalytic components to be assembled<br>(Dockerin+GH) |
| Gene sets independent of<br>cellulosomes |  | B | >=1 | >=0 | >=0 | >=0 | >=1 | microbe-cellulose adhesion<br>(SLH+ CBM) |
|  |  | C | >=0 | >=0 | >=0 | >=1 | >=1 | Cellulose binding cellulase<br>(GH+CBM) |
|  |  | D | >=0 | >=0 | >=0 | >=1 | 0 | Free cellulase<br>(Solitude GH) |

Note: The CBM in this table considered only the cellulose-binding CBM module, and the GH module in this table indicate only the exo/endoglucase GH modules.

Among the carbohydrate active genes independent of the cellulosome gene clusters, what might have been underestimated was the role of genes (type ‘B’) harboring both the SLH module and the cellulose-binding CBM modules. Theoretically, enzymes encoded by these SLH-CBM genes could adhere onto the microbes’ cell surface through its SLH module, and the cellulose-binding CBM counterpart could help drag the SLH-attached microbe cell to its cellulosic substrates. Such microbe-cellulose adhesion facilitated by these SLH-CBM enzymes might help sandwich the excreted carbohydrate enzymes in between the microbe cell and the cellulosic substrate, in which way the CEM complex would form. It is reasonable to speculate that, similar as the hypha mediated CEM complex, the SLH-CBM mediated CEM complex may provide the same physical closeness needed for the synergy among enzymes aggregating in between the microbe cell and the cellulose substrate. There are two other types of cellulosome-independent cellulolytic active genes: type ‘C’ and type ‘D’; both type ‘C’ and type ‘D’ genes harbor the cellulolytic GH modules; and the cellulose-binding CBM modules were identified in type ‘C’ genes but not in type ‘D’ genes.

### Categorization of genomes of potential cellulolytic microbes

As has been summarized in Table S1, among the 2642 microbe strains investigated, only 270 strains were identified with both the exoglucanase GH modules and the endoglucanase GH modules in their genomes. The genomes of these 270 microbe strains harboring both the exoglucanase and endoglucanase GH modules were preliminarily categorized into Group I in this study. Result of the meta-analysis suggested that only genomes in Group I were of potential cellulose hydrolyzing microbes. It was noted that a total number of only one exo/endo GH module (in quite few cases, a total number of two exo/endo GH modules) would be identified in a genome if this genome was annotated with only the exoglucanase GH modules or with only the endoglucanase GH modules, and none of these genomes are of microbe strains with reported cellulolytic activities.

The 270 genomes in Group I could be further categorized into six subgroups (Group I-a, Group I-b, …, Group I-f), according to the types of carbohydrate active genes they harbor. The criteria for this categorization are summarized in Table 2. Cellulosome gene clusters were identified in genomes of the first three subgroups: Group I-a, Group I-b and Group I-c. Unlike that of the Group I-a genomes in which the scaffold genes were of either type A1 or type A2 (with SLH module), the scaffold-genes in genomes of both Group I-b and Group I-c were of type ‘A3’ (without SLH module). The differentiating feature of genomes in Group I-b and Group I-c is that cellulosome-independent SLH-CBM genes were identified in genome of Group I-b, which may act as an alternative microbe-cellulose adhesion machinery; while such cellulosome-independent SLH-CBM genes were absent in genomes of Group I-c.

**Table 2.** Sub-categorization of the genomes in Group I

| Abundance of different gene types identified |  |  |  |  | Category | Number of genomes assigned in each group |
| --- | --- | --- | --- | --- | --- | --- |
| Adhering scaffold (A1 or A2) | Free scaffold (A3) | SLH-CBM (B) | GH_CBM (C) | Solitude GH (D) |  |  |
| $\geq 1$ | $\geq 0$ | $\geq 0$ | $\geq 0$ | $\geq 0$ | Group I-a | 3 |
| 0 | $\geq 1$ | $\geq 1$ | $\geq 1$ | $\geq 1$ | Group I-b | 2 |
| 0 | $\geq 1$ | 0 | $\geq 1$ | $\geq 1$ | Group I-c | 4 |
| 0 | 0 | $\geq 1$ | $\geq 1$ | $\geq 1$ | Group I-d | 16 |
| 0 | 0 | $\geq 1$ | $\geq 1$ | $\geq 0$ | | |
| 0 | 0 | $\geq 1$ | $\geq 0$ | $\geq 1$ | | |
| 0 | 0 | 0 | $\geq 1$ | $\geq 1$ | Group I-e | 139 |
| 0 | 0 | 0 | $\geq 1$ | 0 | | |
| 0 | 0 | 0 | 0 | $\geq 1$ | Group I-f | 106 |

Note: the CBM in this table refers specifically to the cellulose-binding CBM module.

The other three subgroups (Group I-d, Group I-e and Group I-f) were all free of the cellulosome gene clusters. Among these three subgroups, the SLH-CBM genes were identified only in genomes of Group I-d; the cellulose binding CBMs were observed in at least one of the cellulolytic genes in genomes of Group I-e; and genomes of Group I-f were featured with the annotation result that all of their cellulolytic genes were free of the cellulose-binding CBM modules. A detailed summary on the diversity and abundances of the various carbohydrate active genes annotated, and the categorization of these 270 genomes could be found in the Appendix file 2.

### Cellulolytic competency of genomes categorized into the different subgroups

What would the varying genome features indicate on the corresponding microbes’ cellulolytic competency? As has been illustrated in the above section, cellulosome gene clusters are present in three subgroups: Group I-a, Group I-b and Group I-c. Among the 2642 microbe strains, the three strains assigned to Group I-a: *R. thermocellum* ATCC 27405*, R. thermocellum* DSM 1313 and *C. clariflavum* DSM 19732 were all paradigm cellulolytic microorganisms with tethered cellulosome complexes and the highest cellulose hydrolyzing rates reported (24, 26, 27). Both *C. sp.* BNL1100 and *C. cellulolyticum* H10 assigned to Group I-b were reported as proficient cellulose hydrolysers with cellulosome complexes observed (28, 29). There were four strains assigned to Group I-c, being *C. acetobutylicum* ATCC 824, *C. cellulovorans* 743B, *C. acetobutylicum* EA 2018 and *C. acetobutylicum DSM* 1731, respectively; except for *C. cellulovorans* 743B, the three strains of the *C. acetobutylicum* were all inert in crystalline cellulose hydrolysis (8, 28, 30).

Genomes assigned to Group I-d were in two distinct taxonomy groups: strains from the aerobic genus of *Paenibacillus* and strains from the anaerobic genus of *Caldicellulosiruptor*. The seven anaerobic strains in *Caldicellulosiruptor* were all characterized as being cellulolytic (31–37); and the eight aerobic strains of *Paenibacillus* were principally known as plant growth promoter residing either in soil with rich forest residuals or in plant root systems (9, 38–43). The mutualism between *Paneibacillus* and the plant may proceed in a way that the bacteria provide growth hormones and antibiotics to plants, and the plant residues may provide the *Paneibacillus* strains with their carbohydrate substrates.

The total number of the exo/endoglucanase GH modules annotated in genomes of Group I-f varied from 2 to 8, and none of their cellulolytic GH modules were in same genes as the cellulose-binding CBM modules; correspondingly, strains in Group I-f were all inert in cellulose utilization. The total number of the exo/endoglucanase GH modules annotated in genomes of Group I-e were in a wide range of 2-35, and at least one of its carbohydrate active genes harbored both the cellulolytic GH module and the cellulose-binding CBM module. The cellulolytic capacity of microbes in Group I-e varied from being non-cellulolytic to polysaccharides-utilizer to cellulolytic. And there was no apparent correlation between the number of the exo/endoglucanase GH modules annotated and the corresponding microbe’s cellulolytic capability. For example, *Stercorarium subsp.* DSM8532 was cellulolytic with a total number of only 5 exo/endoglucanase GH modules annotated in its genome (44); while *Actinoplanes missouriensis* 431 was not able to grow on cellulose although a total number of 21 exo/endoglucanase GH modules were annotated in its genome (44).

Phylogeny of the 2642 genomes were visualized in the circle tree in Figure 3; genomes assigned into Group I-a, Group I-b, Group I-c, Group I-d and Group I-e were highlighted in different colors; genomes of Group I-f were not highlighted in this circle tree since none of them were cellulolytic. Cellulolytic capability was not highly conservative phylogenetically. For example, among the 13 strains in the genus of *Clostridium* (Appendix file 2), 1 of them was assigned to Group I-b, 4 in Group I-c, 1 in Group I-e; and all the other 7 strains in this genus were not cellulolytic. The results further signified that it might not be a workable approach to predict the corresponding microbe’s cellulolytic capability based solely on the phylogenetic affiliation of a genome.

**Figure 3.**
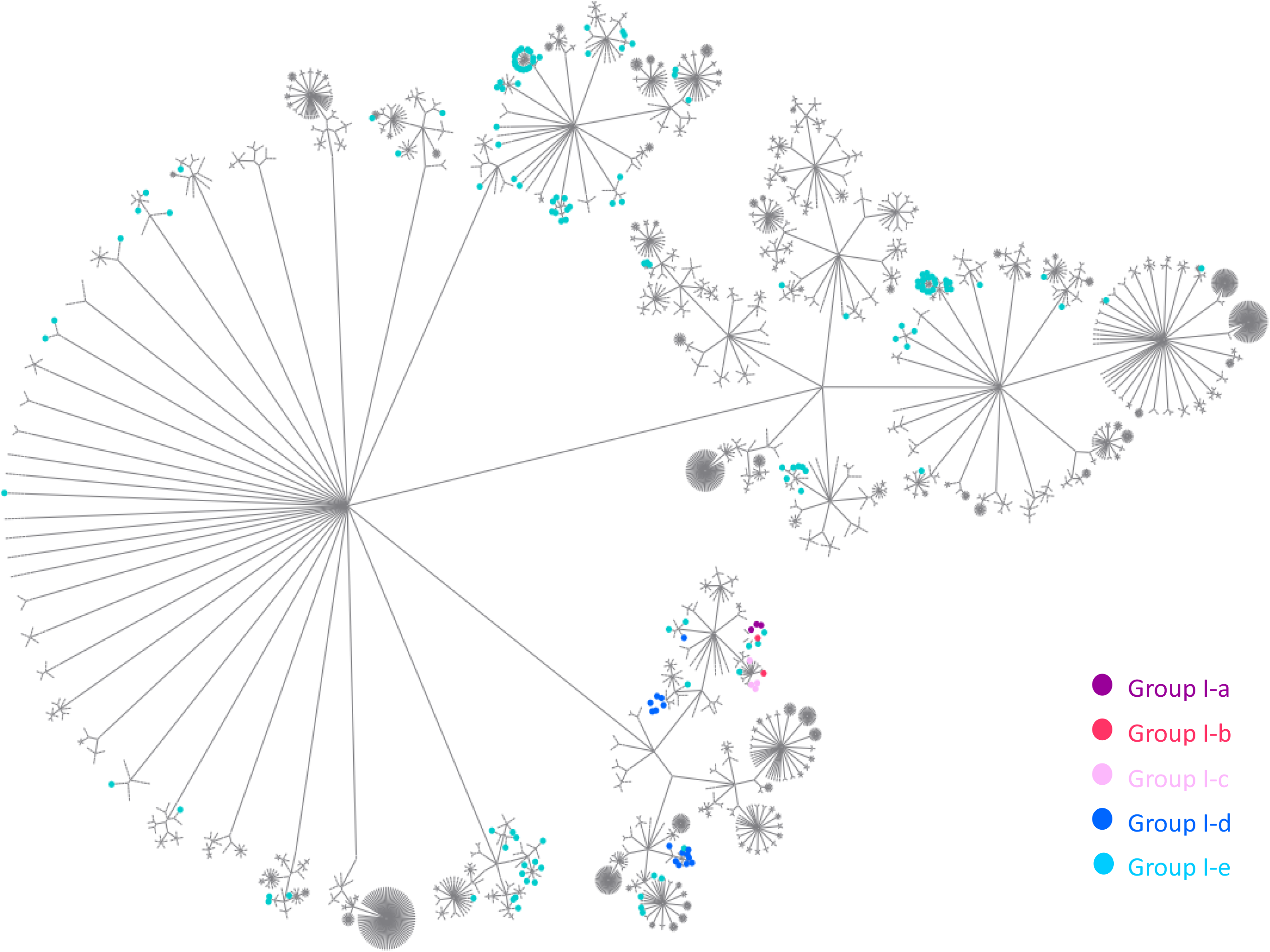
Circle tree of the 2642 genomes. The taxonomy levels of Kingdom (*Bacteria*), phylum, class, order, family, genus and strain were presented by the successive inner nodes. Genomes assigned to the first 5 subgroups of Group I are highlighted in 5 colors.

### The pipeline developed and its application in the annotation of the metagenome assembled genomes on the function niche of cellulose hydrolysis

To facilitate an automatic identification and categorization of the potential cellulolytic genomes, the categorizing criteria proposed in this study were embodied in R scripts. Description of the overall analysis flow and the usage of the scripts could be found in Github. The applicability of this annotation pipeline was further tested with the annotation of the 7904 reference genomes downloaded from NCBI.

Pairing with the dbCAN annotation results, this pipeline was very time-efficient in identifying and categorizing genomes of the potential cellulolytic microbes. It took ~30 minutes to get: 1) a summary of the diversity and abundances of all the CAZy modules identified in each of these 7902 genomes (Appendix file 3); 2) abundances of the diverse carbohydrate active genes in each genome (Appendix file 4); and 3) assignment of the potential cellulose hydrolyzing genomes into 6 subgroups according to the varying synergy machineries annotated (Appendix file 4). Among the 7904 reference genomes annotated, 5 were assigned into Group I-a, 9 genomes were in Group I-b, 15 genomes were in Group I-c, 3 genomes in Group I-d, 15 genomes in Groups I-e and 8 genomes in Group I-f. Figure S1 presents the phylogeny of genomes in the first five sub-groups. Consistent with results of the survey on the 2642 complete genomes, cellulosome-gene clusters were annotated only in a small number of microbes, and the varying cellulolytic capabilities were not phylogenetically conservative.

## Discussion

Cellulosome complexes by its nature could enable the assembly of a number of carbohydrate active units. In previous genome-centric interpretation on the function niche of cellulose hydrolysis, the presence of the cellulosome gene clusters was always taken as an indicator of the efficient cellulose hydrolysers. However, results of this survey suggested that not all cellulosomal gene clusters and the corresponding cellulosome complexes were of the classical configuration, and a finer classification of the cellulosome gene clusters is needed. Cellulosomal complexes in lack of the SLH module might not be cell surface adhering, and the formation of the CEM complex should be aided by some cellulosome-independent SLH-CBM genes. Free cellusomal complexes that could not be held in between the microbe cell and the cellulosic substrate would limit the microbes’ acess to the enzymatic hydrolyzing products, in which case, the host microbe might become relunctant in the energy-consuming synthesis and assembly of cellulosomes. This may explain why the three C. *acetobutylicum* strains were all inert in cellulose hydrolysis although cellulosome gene clusetrs were identified in their genomes.

The potential role of the cellulosome independent SLH-CBM genes in initiating the microbe-cellulose contact was highlighted in this study. Proposed in this study was that the cellulosome independent enzymes encoded by genes (represented by ‘SLH-CBM’ gene in this study) harboring both the SLH module and the cellulose-binding CBM module might be an alternative machinery facilitating the microbe-cellulose adhesion. And such microbe-cellulose contact could further initiate the formation of the CEM complex by sandwiching the carbohydrate active enzymes in between the microbe cells and the cellulosic substrate. The physical closeness in the form of the CEM complex could guarantee: 1) synergy among the enzymes (including the free cellulosome complexes) physically constrained in confined areas, and 2) easy access to the enzymatic hydrolyzing products by the host microbes. C. *acetobutylicum* strains were important for industrial production of acetone/ethanol/proponol, the possibility of C. *acetobutylicum* being able to ferment cellulose would introduce new possibilities for more sustainable solvent production from cheap substrates that include the lignocellulose biomass [48]. One synario proposed in this study to make the C. *acetobutylicum* strains cellulolytic active is by introducing the SLH-CBM genes into their genomes.

One limitation of this study is that we do not think there is no other microbe-cellulose adhesion machineries exist except for the SLH-CBM genes and the cellulosomal complexes, especially in anaerobes. However, the current knowledge on these alternative machineries are limited. For example, glycocalyx containing extracellular polymeric substances (EPS) was reported as a “glue” between the microbe cell and the cellulosic substrates in *R. albus* 7 (45), while we are not sure about the indicator gene for the synthesis of such “glue”. This limitation leads to the uncertainty in the genome-centric interpretation on the cellulolytic capacity of microbes assigned into Group I-c and Group I-e. Novel microbe-cellulose adhesion mechanisms might exist in the cellulolytic microorganisms assigned into Group I-c or Group I-e, e.g., *C. cellulovorans* 743B (Group I-c) and *R. champanellensis* 18P13 (Group I-e). Another factor that needs to be considered in the application of this pipeline is that the quality of the genome matters, more reliable functional interpretation is expected for genomes with higher completeness and lower contamination.

Overall speaking, in the interpretation of MAGs on the function niche of cellulose hydrolysis, the results returned by the annotation approach developed in this study is of good resolution and reliability. Only these genomes assigned into Group I-a, Group I-b, Group I-c, Group I-d and Group I-e are of potential cellulolytic microorganisms. And among these five groups, genomes of Group I-a and Group I-b correspond to cellulolytic microbes with cellulosome complexes. Genomes of Group I-d are of cellulolytic microbes without cellulosome complexes, and the SLH-CBM genes might play essential roles in facilitating the CEM complex formation for microbes in this group. Genomes of Groups I-c and Group I-e might be cellulolytic, while the uncertainty comes not from whether they may harbor potentially novel microbe-cellulose adhesion machineries that could not be recognized by this pipeline.

## Conclusion

In summary, this is the first time that a pipeline was developed for reliable genome-centric interpretation on the function niche of cellulose hydrolysis. The potential cellulose hydrolyzing microbes could be categorized into 5 groups according to the varying synergy mechanisms among the carbohydrate active modules/genes annotated. Pairing with the dbCAN annotation platform, this pipeline is very efficient in identifying potential cellulose hydrolysers by interpreting the complete genomes or MAGs recovered through high-throughput sequencing.

## Methods

5243 GenBank Format (GBK) files corresponding to 2786 prokaryote with complete genomes were downloaded from the NCBI genomes_Ftp_Site (ftp://ftp.ncbi.nlm.nih.gov/genomes/archive/old_genbank/Bacteria/). The reason why this old archive collection (last updated on Dec. 2^nd^, 2015) was chosen in this study was that, comparing with the most recently updated achieve, this collection had a higher portion of complete genomes from strains whose phenotypes have been well characterized; and the documented phenotypes make it possible to evaluate the reliability of the genome-centric prediction on the corresponding microbes’ cellulolytic capability. Another batch of 7904 reference genomes were also downloaded from NCBI (ftp://ftp.ncbi.nlm.nih.gov/genomes/refseq/bacteria/) (updated on February, 2019), and there are metagenome assembled genomes (MAGs) among these 7904 reference genomes. These 7904 reference genomes were used to evaluate the applicability of the pipeline in the batch annotation of a large number of MAGs. A detailed summary on these 7904 reference genomes could be found in Appendix file 5.

Fasta Amino Acid sequences (FAA) of the coding regions (often abbreviated as CDS) were extracted from the GBK files with a python script. The FAA files were then subjected to the dbCAN HMMsearch for the CAZy module annotation, following the HMMsearch criteria (e.g. cutoff value) recommended by the dbCAN developers (6). CAZy (carbohydrate active enzymes) modules were identified in 3898 of these FAA files that corresponded to 2642 prokaryotic strains. The assembly accession numbers and taxonomy affiliation of these 2642 strains have been summarized in the Appendix file 6. As the chromosome and the plasmid in one same microbe strain have separate FAA files, results of the annotation of those separate FAA files of the chromosome and the plasmid in one same microbe strain would be aggregated to represent all the CAZy modules annotated in one microbe strain.

The GH modules that were relevant in the cellulose hydrolysis were classified and read as the exoglucanase GH modules, the endoglucanase GH modules, the xylanase GH modules and the glucosidase GH modules, respectively (Table S2). The CBM modules were classified and read as the cellulose-binding CBM modules, the hemicellulose-binding CBM modules and other CBM modules, respectively (Table S3). The dockerin, cohesion and the SLH modules were the three important accessory modules in the cellulosome gene clusters. Based on the survey of the carbohydrate active genes in the 2642 complete genomes, frequencies of the CAZy modules co-occurring with one another in same genes were calculated; and the principles applied in such calculation could be found in the supporting information.

Applying the genoplotR package in R (46), the CAZy module arrangement along genes in genomes could be visualized. Batch visualization of the arrangement of the CAZy modules along all the carbohydrate active genes annotated in each complete genome or MAG could be achieved. Scripts of the pipeline and workflow of the pipeline have been well documented in Github. In addition to the interpretation of the complete genomes from the 2642 CAZy-harboring strains, and the 7904 reference genomes downloaded from NCBI, the pipeline developed in this study was also applied in the annotation of 17 metagenome assembled genomes (MAGs) recovered from a cellulose converting consortia enriched in our previous study (47). These 17 MAGs can be applied as an example dataset to work with, and all the raw data and results generated on these 17 MAGs have also been deposited in the Github.

## Availability of data and materials

All data generated or analyzed during this study are included in this manuscript, its supplementary information files and the appendix files. All scripts written in this study are available in https://github.com/yuboer/genome-centric-portrait-of-cellulose-hydrolysis.

## Additional files

Appendix file1: CAZy modules cooccurring frequencies along genes

Appendix file2: Further categorization of complete genomes harboring both the exoglucanse and endoglucanase GH modules

Appendix file3: Abundance and diversity of CAZy modules in each of the 2642 complete genomes

Appendix file4: Summary of the carbohydrate active genes annotated and categorization of the 270 complete genomes in Group I

Appendix file5: NCBI accession of the 7904 MAGs investigated

Appendix file6: Accession numbers of the 2642 complete genomes investiagted

## Abbreviations

SLH: Surface Layer Homology
CBM: Carbohydrate Binding Module
CEM: Cellulose-Enzyme-Microbe
MAGs: Metagenome Assembled Genomes (MAGs)
NGS: Next Generation Sequencing
CAZy: Carbohydrate Active enzyme
PUL: Polysaccharide-Utilization Loci
EPS: Extracellular Polymeric Substances

## Authors’ contributions

Yubo Wang conceived the study, analyzed the data and wrote the manuscript. Liguan Li contributed resources of the 2642 complete genomes and the corresponding metadata collection; Yu Xia contributed in the CAZy modules annotation. Feng Ju contributed by providing constructive suggestions during the writing of this manuscript. Tong Zhang conceived the study. All authors edited the manuscript and approved the final draft.

## Acknowledgments

Yubo Wang wish to thank the University of Hong Kong for the postgraduate scholarship. Liguan Li and Yu Xia acknowledge the postdoc scholarship provided by the University of Hong Kong.

